# Biosynthesis of linear protein nanoarrays using the flagellar axoneme

**DOI:** 10.1101/2021.09.02.458790

**Authors:** Hiroaki Ishikawa, Jie L. Tian, Jefer E. Yu, Wallace F. Marshall, Hongmin Qin

## Abstract

Applications in biotechnology and synthetic biology often make use of soluble proteins, but there are many potential advantages to anchoring enzymes to a stable substrate, including stability and the possibility for substrate channeling. To avoid the necessity of protein purification and chemical immobilization, there has been growing interest in bio-assembly of protein-containing nanoparticles, exploiting the self-assembly of viral capsid proteins or other proteins that form polyhedral structures. But these nanoparticle are limited in size which constrains the packaging and the accessibility of the proteins. The axoneme, the insoluble protein core of the eukaryotic flagellum or cilium, is a highly ordered protein structure that can be several microns in length, orders of magnitude larger than other types of nanoparticles. We show that when proteins of interest are fused to specific axonemal proteins and expressed in living cells, they become incorporated into linear arrays which have the advantages of high protein loading capacity, high stability, and single-step purification with retention of biomass. The arrays can be isolated as membrane enclosed vesicle or as exposed protein arrays. The approach is demonstrated for both fluorescent proteins and enzymes, and in the latter case it is found that incorporation into axoneme arrays provides increased stability for the enzyme.

## Introduction

We often contrast biological systems with manmade systems by viewing biological machines as squishy and stochastic, compared to manmade machines that are rigid and precisely organized. This difference creates challenges for engineering biological systems to perform defined functions. As a result, much of biotechnology has historically focused on solution-phase enzyme systems. When it becomes necessary to immobilize enzymes or assemble them into structured arrays, this is done by isolating the enzymes and then using various physical or chemical means to immobilize them on artificial surfaces or particles. In certain cases, however, biology is able to build highly sophisticated and complex assemblies. Indeed, Drexler has proposed that biological systems exist as a distinct phase of matter he calls the “machine-phase” (Drexler, 1992). One of the most extreme examples is the eukaryotic flagellar axoneme, a linear structure in which hundreds of different proteins are docked in precise positions and defined orientations. We propose that the highly organized and regularly arranged structure of the axoneme can be engineered to allow stoichiometric production of protein components as immobilized linear nanoarrays. This approach confers several advantages, such as high protein loading capacity compared to other bioparticle systems; genetically programmed self-assembly without the need for any crosslinking steps; single-step purification of particles without the need for cell lysis, allowing retention and re-use of biomass; and choice of isolating the particle as a membrane enclosed vesicle or as an exposed protein array. This work will serve as the foundation for a new approach to machine-phase synthetic biology that harnesses the precise assembly of cytoskeletal structures.

One straightforward application of solid-phase synthetic biology is in biocatalyst enzyme production, a core methodology in biotechnology. With the desire for increased protein production, multiple biotechnological methods have been developed. Enzymes are typically expressed in bacteria (Rosano et al., 2019), yeast (Baghban et al., 2019), insects, or mammalian cells (McKenzie and Abbott, 2018) using recombinant molecular approaches. Optimizing these systems can require substantial effort to solubilize and purify the enzyme of interest. Moreover, even after enzymes are successfully produced and purified, they are easily degraded or inactivated, and difficult to store. Enzyme immobilization is an effective strategy to maximize both the physical and the enzymatic stability of a biocatalyst (Basso and Serban, 2019; Tufvesson et al., 2010).

Immobilization strategies have evolved from simple physical absorption or covalent cross-linking onto insoluble carriers, such as agarose and Sepharose beads, or polypropylene, to the current focus on encapsulating enzymes into nanoparticles (Azuma et al., 2016; Datta et al., 2013; Hagen et al., 2018; Plegaria and Kerfeld, 2018). Encapsulation can provide stability and protection from the environment. The local concentration of protein within a particle can be extremely high, potentially allowing channeling of reaction products if enzymes responsible for sequential steps are included together in the same particle. This channeling can lead to more efficient overall reactions as well as restricting the release of potentially toxic intermediates (Giessen and Silver, 2016; Khattak et al., 2014). These desirable traits on medicine, research, and biotechnology have provided the impetus to search for systems and engineering approaches suitable for protein encapsulation. However, a major challenge remains how to incorporate a purified enzyme into a nanoparticle, via crosslinking or other strategies for attachment.

Naturally occurring self-assembled protein nanoparticles are excellent candidates for engineering protein encapsulates. For example, the carboxysome responsible for carbon fixation in some bacteria (Rae et al., 2013) and the polyketide synthase particle (Fischbach and Walsh, 2006) can be harnessed to build synthetic nanoparticles containing a virtually limitless range of possible enzymes or other proteins of value. Nanoreactors based on bacteriophage MS2 capsid proteins (Giessen and Silver, 2016) and the non-viral lumazine synthase protein (Azuma et al., 2016) have been experimentally tested for their ability to encapsulate fixed quantities of enzymes. Both proteins assemble into polyhedral shapes of defined size, creating a capsule with a lumen into which suitably tagged enzymes can be docked during particle assembly via protein-protein interactions. One limitation is the quantity of protein that can be incorporated. This is because the polyhedral geometry of the particle imposes a strict size limitation. Another limitation of these tightly packaged protein aggregates is that the enzyme’s activity could be adversely affected due to misfolding or trapping toxic substrates (Azuma et al., 2016)

Linear protein arrays present one possible alternative structure in which the size, and therefore the quantity of protein incorporated, could be much greater than in smaller polyhedral particles. There are many examples of proteins that self-organize into linear arrays within the cell. Examples include cytoskeletal filaments such as actin and tubulin as well as enzyme filaments such as glutamate synthase, CTP synthase, and the eIF2/2B complex (Noree et al., 2010). While self-assembling filaments offer a way to increase protein content, they suffer from a much higher variability in total quantity. A population of self-assembling linear polymers is predicted to show an exponential distribution of lengths (Oosawa and Kasai, 1962), a prediction that has been confirmed through analysis of length distribution of actin and other polymers assembled in vitro (Kawamura and Maruyama, 1970). Constraining the polymerization to the interior of a cell can narrow the length distribution somewhat (Gregoretti et al., 2006; Janulevicius et al., 2006). Nevertheless, intracellular polymers show wide length distributions of at least several fold, making them non-ideal as protein carrier nanoarrays unless some method can be used to constrain length variation. While several naturally occurring self-assembled protein nanoparticles and linear systems are available for engineering protein encapsulates, they are limited to particles with a diameter of 10 to 150 nm (Lopez-Sagaseta et al., 2016) or are difficult to implement due to intrinsic system variability. Currently, no protein expression platform in use is capable of directly self-assembling protein nanoarrays with defined size in living cells.

The flagellar axoneme of the green alga *Chlamydomonas reinhardtii* presents an ideal scaffold for constructing linear protein nanoarrays (**Figure 1**). The eukaryotic flagellum is a motile structure that consists of a protrusion of the plasma membrane supported by the axoneme, a protein-based assembly consisting of nine doublet microtubules (DMTs) together with several hundred associated proteins involved in building the axoneme and driving flagellar motility. Flagellar proteins are synthesized in the cell body first and then make their way to the flagellum. A complex molecular machinery known as the intraflagellar transport (IFT) system uses a combination of motors and protein chaperones to transport insoluble proteins into the axoneme and incorporate them into the appropriate positions (Ishikawa and Marshall, 2011; Qin, 2012; Qin et al., 2004). Many different proteins incorporate into the axoneme with fixed spatial repeats, for example, radial spokes and dynein arms binds to the axoneme with an underlying 96 nm periodicity that is generated by molecular rulers aligned to the axonemal lattice (Oda et al., 2014). The *Chlamydomonas* axoneme therefore provides an excellent platform for the self-assembly of proteins on DMTs with high density and precise periodicity (**Figure 1A**).

**Figure 1.**
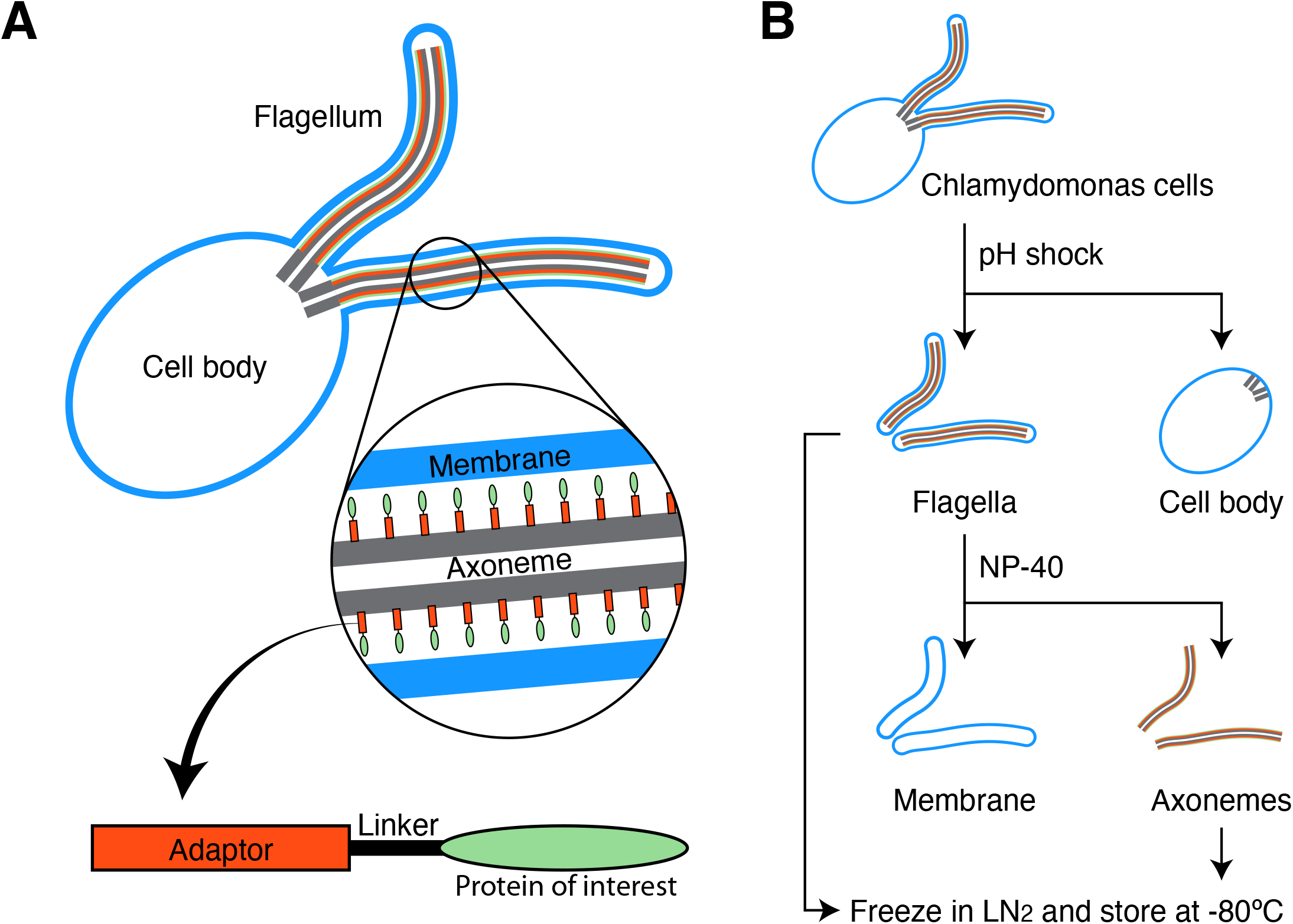
Harnessing the flagellar axoneme as a biologically self-assembling protein nanoarray. (A) *Chlamydomonas* algal cells contain two flagella that protrude from the cell surface. The structural core of the flagellum is the axoneme, an array of nine doublet microtubules that is, in turn, surrounded by the flagellar membrane. By fusing a protein of interest to one of the axonemal proteins, the axoneme protein can serve as an adaptor to attach many copies of the protein of interest into the axoneme, forming a protein array. (B) One-step purification and low-cost harvesting of protein arrays in multiple forms. Flagella can be cleanly detached from the cell body by transiently reducing the pH of the media (pH shock), which releases the cell body intact, allowing the flagellum to be isolated in a single centrifugation or filtration step. The flagellum can be stored in this form, or can be treated with detergent to remove the membrane, producing a naked axoneme where the fusion proteins are directly exposed. This system therefore allows the protein array to be isolated in either a membrane-bound vesicle form or a solvent exposed membrane-less form.

In contrast to small sized polyhedral shape nanoparticles, axonemes typically are several orders of magnitude larger, on the 1-10 micron length scale. But unlike other large linear arrays such as microtubules or actin filaments, axonemes are subject to narrow length variation, typically showing a sharper-than-Gaussian (Kannegaard et al., 2014) length distribution. Importantly, the length of the axoneme can be tuned using a collection of existing mutations (Silflow and Lefebvre, 2001), including mutants that have flagella that are longer than normal (Asleson and Lefebvre, 1998; Barsel et al., 1988; McVittie, 1972), and other mutants that have shorter flagella (Ishikawa et al., 2014; Kuchka and Jarvik, 1987). Length can also be tuned using chemical inhibitors. For example, lithium causes flagella to increase length (Nakamura et al., 1978), while other compounds obtained in chemical screens cause flagella to shorten (Avasthi et al., 2012; Engel et al., 2011). Flagellar length is thus tunable via both genetic and chemical means.

Compared to other types of biological nanoparticles, axonemes are extremely easy to purify (**Figure 1B**). Flagella can be detached from living cells with a simple pH drop and purified with a single centrifugation step (Witman et al., 1972). The cell bodies remain intact and viable, allowing for virtually complete recovery of biomass for further rounds of protein array production. The axoneme can then be obtained by treating the purified flagella with detergent, or else the whole flagellum can be used as a membrane-encapsulated array. Moreover, isolated flagella are stable and easily stored because no protease exists in the flagellar compartment (Pazour et al., 2005).

As a source of biological material, the unicellular green alga *Chlamydomonas reinhardtii*, a genetically tractable green alga, can be grown at industrial scale using inexpensive media (Silflow and Lefebvre, 2001), and has been considered as a potential organism for production of algal biofuel (U.S. DOE 2010). It can be cultured in large quantities with or without light. Other features that make this organism particularly useful include that it is safe for human consumption and flagella can be repeatedly regenerated and isolated. Upon removal of a flagellar axoneme, *Chlamydomonas* will regrow the structure within two hours (Rosenbaum et al., 1969), thus promoting rapid synthesis of desired protein products in a uniform array. Here we describe the use of the flagellar axoneme from *Chlamydomonas reinhardtii* as a platform for synthesis of self-assembled protein arrays in living cells.

## Results and Discussion

### Evaluating axonemal proteins as suitable adaptor for enzymes

The axoneme is a complex structure containing hundreds of different proteins (Pazour et al., 2005). In principle any of these could be used as an adaptor by fusing a protein domain of interest, such as an enzyme, onto one end of the axonemal protein (**Figure 1A**). Then the adaptor fusion construct could be expressed in mutant cells that lack the endogenous copy of the adaptor protein encoding gene

As proof of concept, we examined three candidate adaptor proteins: IFT20, RSP3, and FAP20 (**Figure 2**). IFT20 is a protein subunit of IFT particles (Cole et al., 1998). RSP3 is part of the radial spoke complex, a T-shaped complex that is composed of at least 23 proteins (Curry and Rosenbaum, 1993). The radial spoke stalk is anchored to the inner side of doublet microtubules and its globular head extends toward the central pair of singlet microtubules. In each 96 nm periodic structure, there are two complete radial spokes and one truncated spoke (Yang et al., 2006). RSP3 is encoded by the *PF14* gene, and *pf14* loss-of-function mutants have paralyzed flagella (Luck et al., 1977). FAP20 is a component of the inner junction (IJ) filament complex, a protein complex located at the seam between the A and B tubules of the outer doublet microtubules (Khalifa et al., 2020). FAP20 lines the IJ of the doublet microtubule at a precise 8 nm periodicity, forming 9 parallel solid nanoarrays (Yanagisawa et al., 2014). The relative locations of these three adaptor proteins are shown in **Figure 2A**.

**Figure 2.**
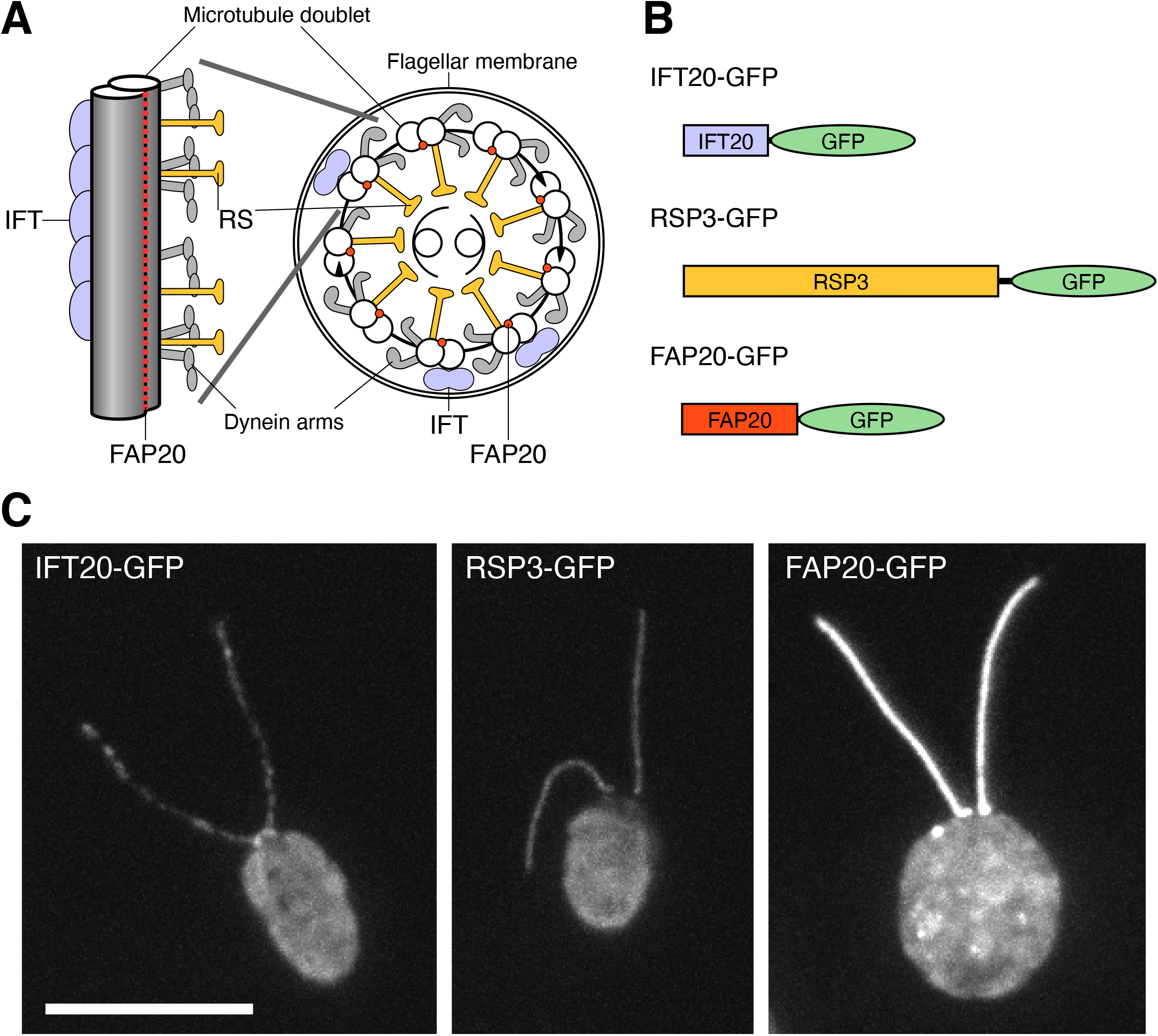
Adaptor proteins for axonemal nanoarray. (A) Cross section and longitudinal views of axoneme showing location of IFT20, RSP3, and FAP20 in their respective complexes. RS, radial spoke; IFT, intraflagellar transport. (B) Diagram of constructs showing where GFP is fused to the protein. (C) *Chlamydomonas* cells expressing GFP-tagged fusion constructs with IFT20, RSP3, and FAP20. Scale bar, 10 μm.

To evaluate adaptor proteins, we expressed GFP-tagged versions of these three candidate adaptor proteins. Fusions of IFT20 (Follit et al., 2006), RSP3 (Diener et al., 1993), and FAP20 (Yanagisawa et al., 2014) (**Figure 2B**) have all been previously described in the literature. Importantly, all three were shown to rescue loss of function mutations in the corresponding gene even when fused to GFP, indicating that the GFP fusion does not interfere with function. Visual comparison (**Figure 2C**) showed that FAP20-GFP showed the brightest signal, indicating the largest quantity of protein incorporated into the axoneme.

As a quantitative comparison for incorporation among the different scaffold proteins, we implemented an automated segmentation procedure for flagella, which traces the linear background of each flagellum, and quantifies fluorescence intensity as a function of position (**Figure 3A**). Segmentation of flagella was performed by first identifying the cell body, subtracting it from the image, and then traversing each flagellum, starting at the two tips and identifying neighboring pixels of maximal intensity. This method was able to reliably label each flagellum (**Figure 3B**) allowing GFP intensity to be quantified specifically inside the flagella, without interference from the cell body. Using this algorithm, we quantified the total fluorescence intensity in each of the three strains (**Figure 3C**). In all three cases, we found that the total intensity scales linearly with the flagellar length. For any particular length, FAP20-GFP always gave the highest total intensity, and RSP3 the lowest, suggesting that FAP20 scaffold is more effective than the other two in terms of protein incorporation into the flagellum. Scatter of points around the best fit lines in **Figure 3C** indicated that IFT20 shows substantially greater variation than RSP3 which had a similar average intensity. This is perhaps to be expected, given that the IFT complexes localize sparsely along the flagellum and undergo constant transport in and out of the flagellar compartment (Ludington et al., 2015). This analysis demonstrates that for all constructs tested, incorporation is directly proportional to length, but that compared to IFT20 and RSP3, FAP20 shows the highest level of incorporation per unit length. We also noted visually in **Figure 2** that FAP20 shows the most spatially uniform incorporation along the length of the flagellum. These results indicate that FAP20 may be the most suitable scaffold protein for building axonemal protein arrays.

**Figure 3.**
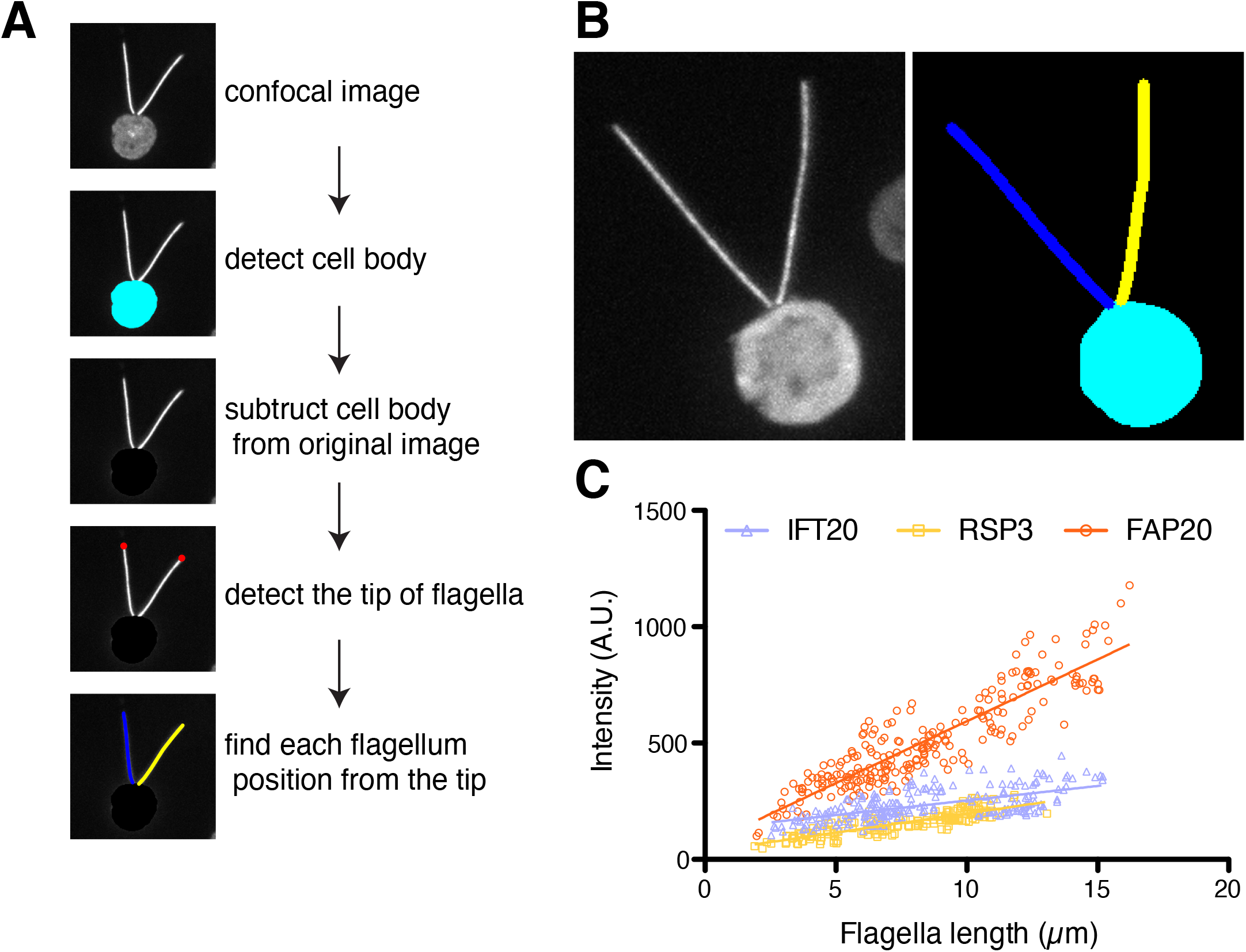
Quantification of GFP incorporation into flagella. (A) Diagram to show the steps of the image analysis algorithm. (B) illustrative example showing segmentation of the flagella. (C) Graph of total GFP intensity versus flagellar length for the three constructs.

### Stability of protein arrays during isolation and storage

Easy isolation and high stability during both isolation and storage are essential for any sort of nanoparticle or nanoarray to be useful in practice. *Chlamydomonas* cells sever flagella at the base when placed under stress conditions, for example pH shock (Quarmby and Hartzell, 1994). The flagella can then be cleanly separated away from the cell bodies by centrifugation through a sucrose cushion (Richey and Qin, 2013). These flagella can then be frozen down for storage or else demembranated to produce isolated axonemes (**Figure 1B**).

Axonemes are stable structures, but the degree to which individual axonemal proteins can remain stably anchored to axonemes has never been systematically addressed. We have found that FAP20-GFP is retained during isolation of flagella as well as following demembranation to produce axonemes (**Figure 4**). For both flagella and axonemes, FAP20-GFP was retained during freezing and thawing (**Figure 4E**). The high stability of protein arrays on flagellar axoneme during isolation and storage is desirable for practical usage.

**Figure 4.**
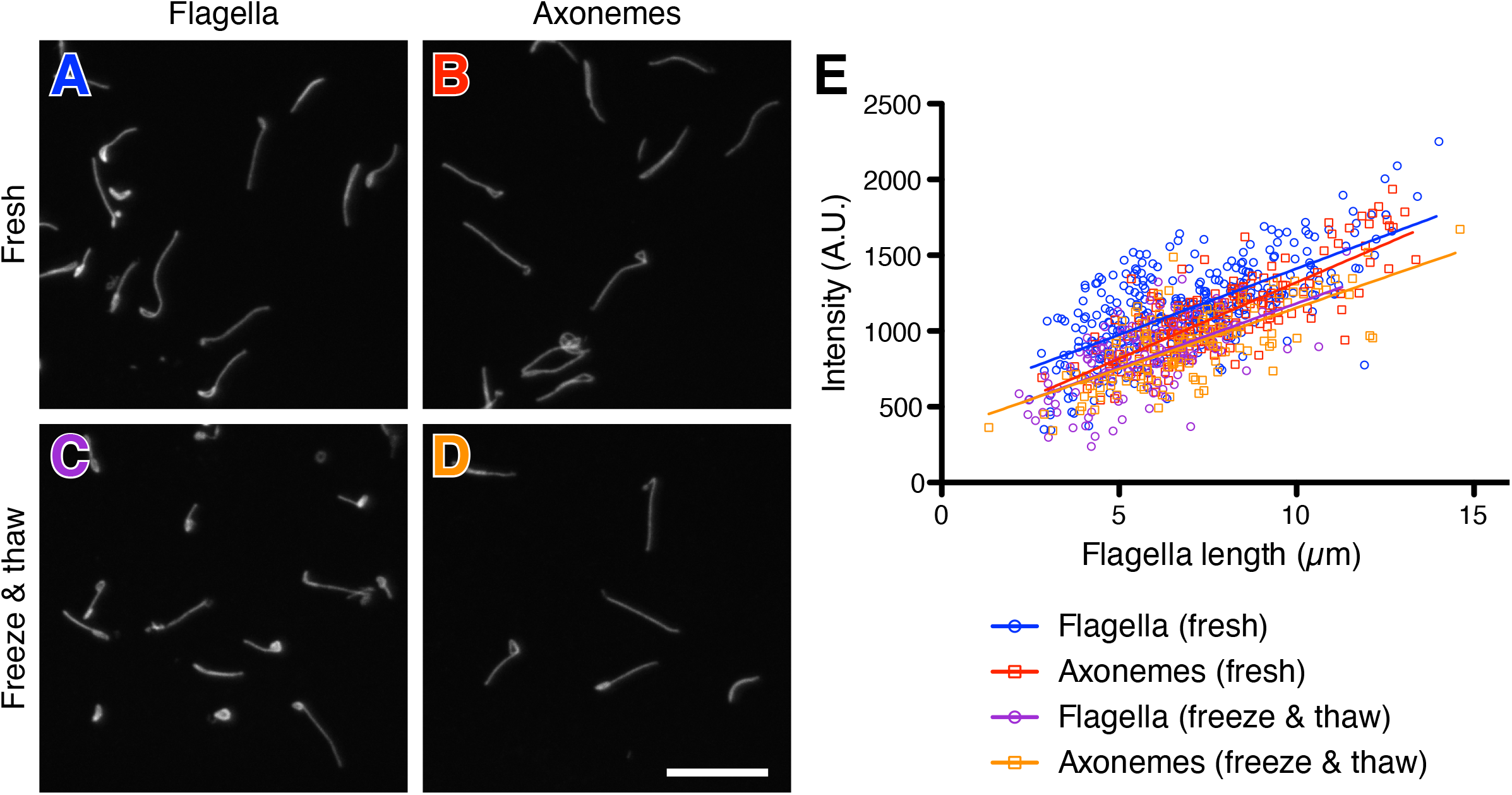
Stability of FAP20-GFP during nanoarray isolation and storage. (A,B) Images of detached flagella following pH shock before (A) and after (B) demembranation to produce axonemes. (C,D) Detached flagella (C) and axonemes (D) following storage at -80°C. Scale bar for (A-D) 10 μm. (E) Quantification of GFP fluorescence showing good retention of signal during axoneme preparation. Freeze-thaw does result in a small decrease in signal.

### Using β-Lactamase to evaluate biosynthesis of enzyme axonemal nanoarrays

As a proof of concept for enzyme incorporation into axonemal arrays, we expressed β-Lactamase (βLac) fused to FAP20 and RSP3 as adaptors to evaluate if flagellar axonemes are suitable for biosynthesis of enzyme nanoarrays. βLac has been tested in self-encapsulation using lumazine synthase protein (Azuma et al., 2016), in which it could be efficiently encapsulated, but the encapsulated enzyme showed more than a 10-fold drop in catalytic efficiency compared to soluble βLac. Protein misfolding was likely responsible for the reported loss of the catalytic activity, which is notoriously difficult to deal with. We expressed FAP20-TEV-βLac or RSP3-TEV-βLac construct (**Figure 5A**) in the non-motile loss-of-function mutant *im5 (fap20)* or *pf14 (rsp3)* (Yanagisawa et al., 2014), respectively. The cells expressing the fusion proteins became motile because of the expression of adaptor proteins and were identified by microscopic observation (**Supplemental Movies S1-S5**). The yield of products was accumulated by flagellar regeneration. FAP20-βLac flagella were regenerated once, and RSP3-βLac flagella were regenerated twice due to the shorter flagellar regeneration time. The protein concentration of all flagella samples was quantified by comparing to BSA standard samples (**Supplemental Figure S1**). The concentration of each sample was adjusted to 1.6 μg/μl and then subjected to Western blotting. The expression of target protein βLac was immunoblotted by βLac antibody, wild-type (WT) flagella samples were used as a negative control. Immunoblotting of acetylated tubulin confirmed that all samples had equal loading (**Figure 5B**).

**Figure 5.**
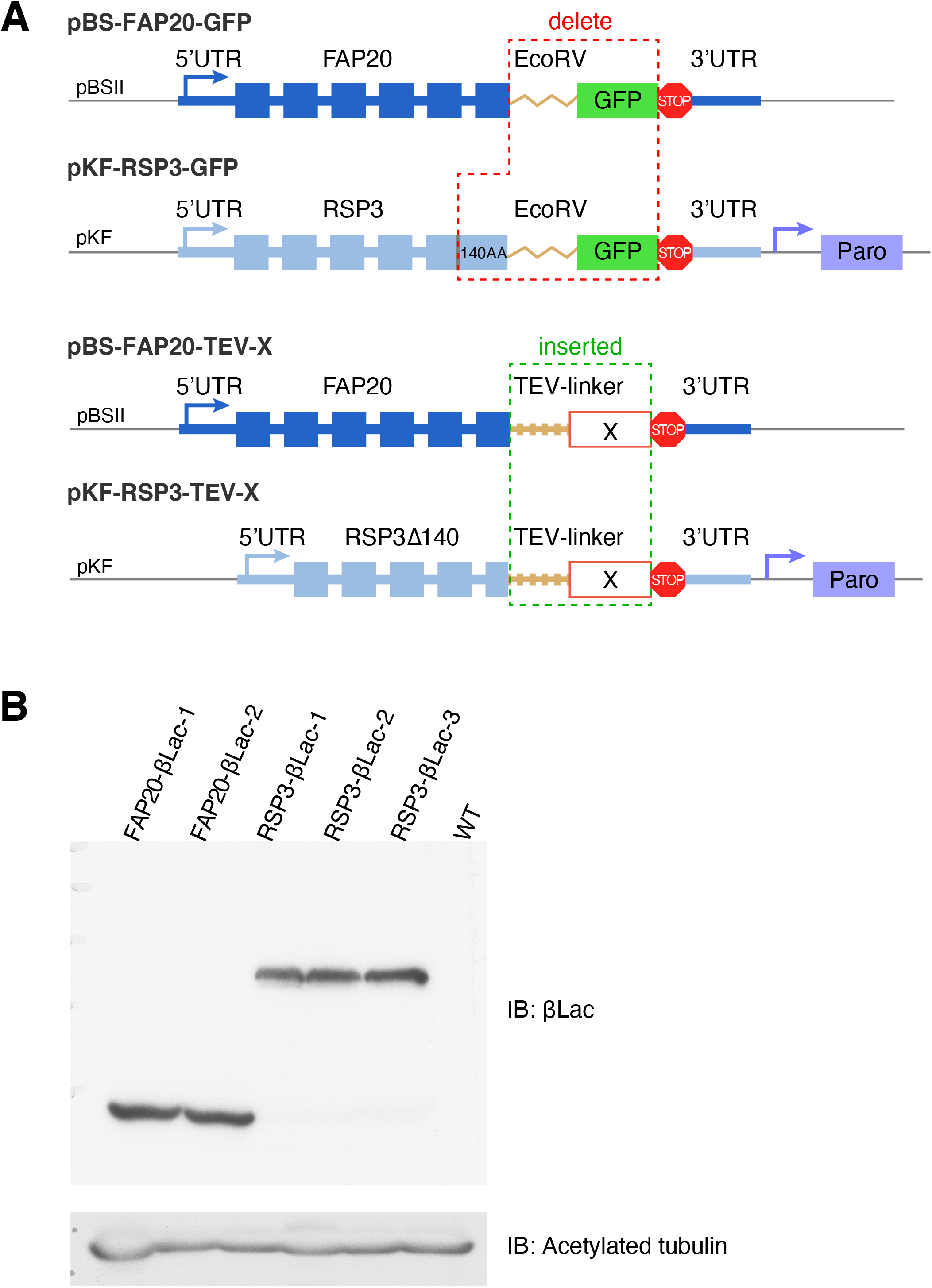
Constructs for expressing fusion proteins. (A) The constructs of pBS-FAP20-GFP and pKF-RSP3-GFP (Yanagisawa et al., 2014) were modified to express the fusion proteins. The fusion protein was connected to FAP20 or RSP3 with a TEV-linker. The restriction enzyme EcoRV digestion sequence and the GFP gene were deleted in both of the original constructs. In addition, the C-terminal 140 amino acids of RSP3 were deleted as well in the pKF-RSP3-GFP construct. The TEV-linker and the target gene (X), such as βLac, were fused to the C-terminus of either FAP20 or RSP3. (B) The concentration of flagella samples was adjusted to 1.6 μg/μl with the dilution of HMDEK buffer and subjected to Western blotting. The immunoblotting of βLac showed the protein expression levels in each of the flagella samples and immunoblotting of acetylated tubulin showed the equal loading of each sample.

We measured the βLac catalytic activity using a colorimetric Beta Lactamase Activity Assay Kit (**Figure 6** see Methods) to compare the catalytic efficiency of FAP20-TEV-βLac or RSP3-TEV-βLac in the isolated flagella to commercial βLac enzyme. Wild-type *Chlamydomonas* cells do not possess measurable endogenous βLac activity, but when either the FAP20 or RSP3 fusions with βLac were expressed, isolated flagella showed clear enzymatic activity that was proportional to the quantity of flagella tested (**Figure 6A**). Incorporation into the axoneme was apparently not deleterious for enzyme activity. In fact, when the same amounts of protein were used, the catalytic efficiency of βLac anchored on the axoneme was about six-fold higher than that of soluble βLac (**Figure 6B**), indicating that βLac enzyme is correctly folded when assembled on the axoneme. Furthermore, the fixed enzymes on axonemes appeared more resistant to freeze-thaw treatment, and more stable for room temperature storage. Protein arrays also retained enzymatic activity when separated from substrate and then used for a new round of assays (**Figure 6C**).

**Figure 6.**
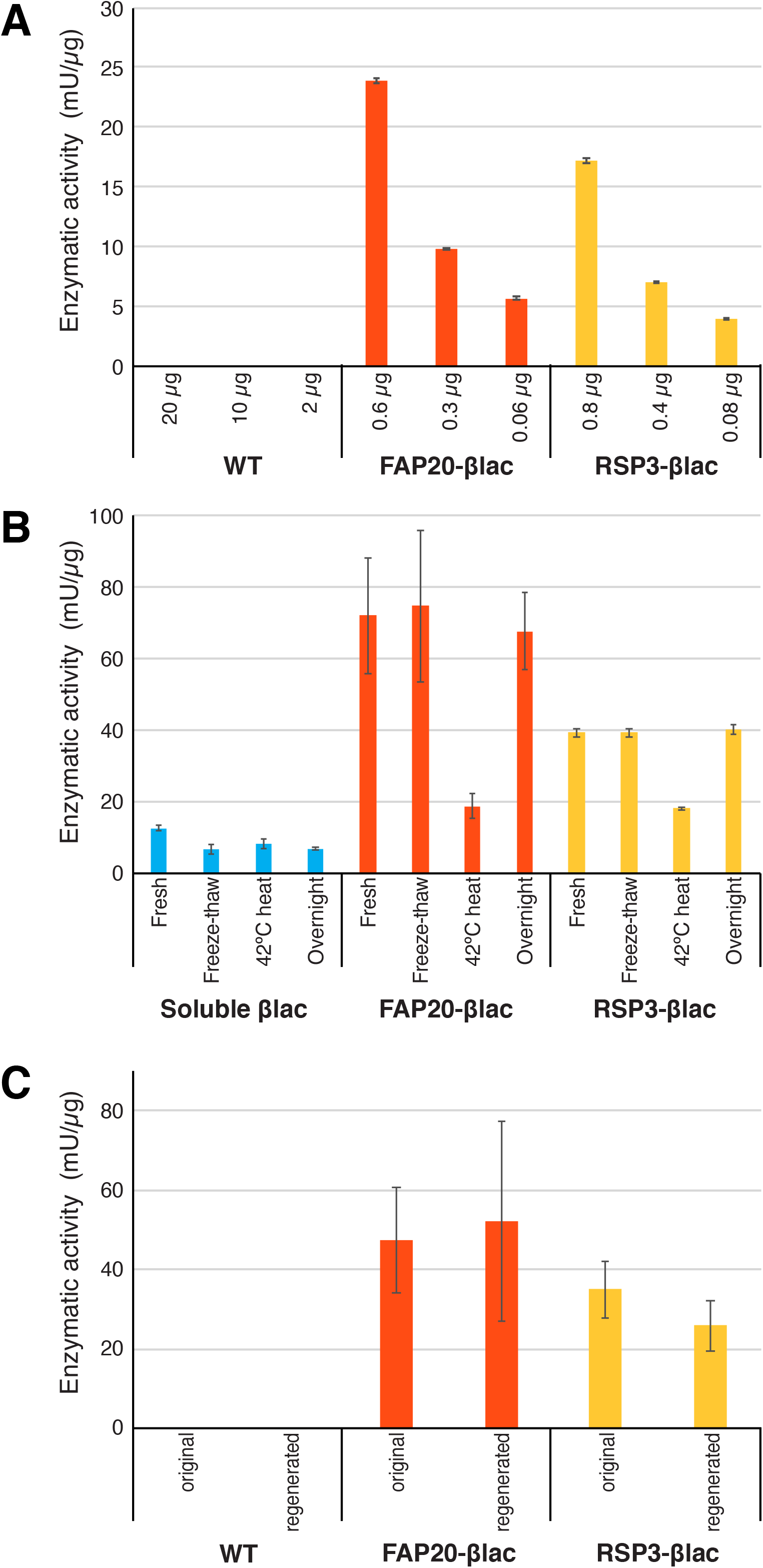
Axoneme-based enzyme arrays using βLac. (A) βLac activity is undetectable in WT *Chlamydomonas* flagella but is conferred by flagella containing FAP20 or RSP3 βLac fusion constructs. (B) βLac activity in axonemes compared with soluble enzyme. (C) Recovery of enzyme activity in recycled flagella. The flagella from the first reaction were collected and re-subjected to a new reaction. Two replicates were performed for the enzymatic activity assay.

### Release of attached proteins by TEV protease cleavage

We envision the axonemal array system as a flexible platform for building structured arrays of immobilized proteins in fixed relative orientations. However, because the IFT system can handle insoluble proteins as part of its normal function, we propose that axonemal protein arrays might also be useful as protein expression systems. In this case, a method would be needed to release the protein from the rest of the axoneme. To test this idea, we generated a variant construct in which the FAP20 and GFP domains were linked by a linker containing a TEV protease cleavage site (**Figure 7A**). Flagella were harvested from strains expressing this construct, and used to prepare axonemes. Experiments confirm that the vast majority of GFP was removed following TEV protease treatment of the isolated axonemes (**Figure 7B**).

**Figure 7.**
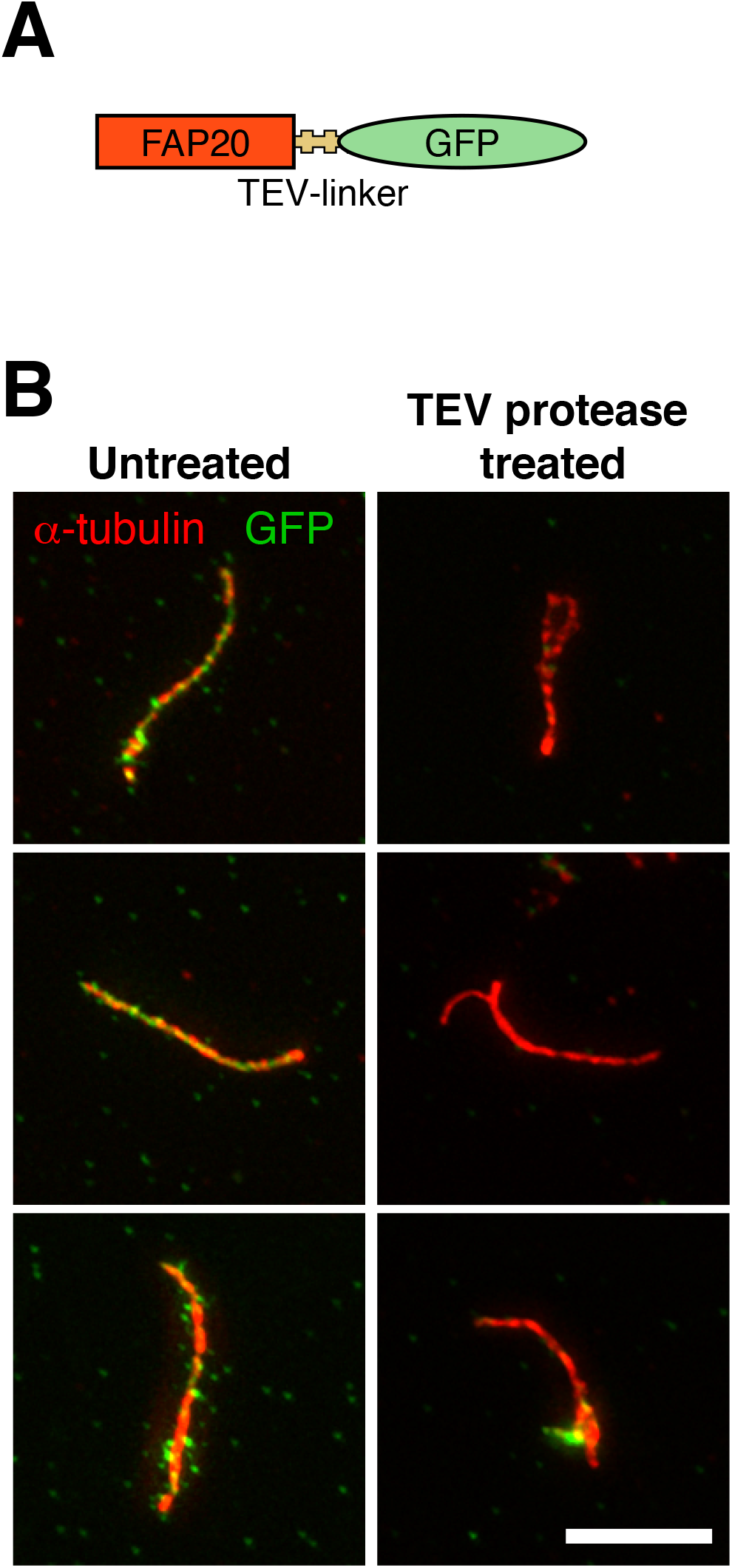
Release of fusion constructs by TEV protease treatment. (A) Diagram of modified construct that includes a TEV protease cleavage site. (B) Bottom panels are three examples of TEV treatments. Comparing untreated versus protease treated axonemes, in all cases fixed and stained to detect acetylated tubulin (red) and GFP (green). Note the granular appearance of the GFP signal compared to earlier images is an artifact of methanol fixation required to image acetylated tubulin. Reduced green fluorescence in axonemes treated with TEV protease demonstrates release of most of the GFP domain from the fusion construct. Scale bar, 5 µm.

### Design challenges for axonemal enzyme arrays

We have shown that the axoneme is a flexible platform for biological assembly of protein arrays, able to incorporate at least two structurally distinct protein clients (GFP and βLac) while retaining their function. Future work will be needed to determine the range of proteins that are compatible with this platform. The size, charge, and quaternary structure of target proteins may affect their ability to enter the flagellar compartment as well as to fit stably into the context of the axoneme. These constraints are not unique to axonemes - in other cases of biological nanoparticles, protein charge and size clearly affects the level of incorporation (Azuma et al., 2016). The evolutionary selection of the IFT system to transport extremely large protein complexes such as dynein arms and radial spokes suggests that axonemal assembly may be particularly amenable to a wide range of target proteins.

Given that an enzyme can be efficiently incorporated into an axonemal array, the next concern is to ensure that the enzyme will be active. Enzymes can be strongly affected by their chemical environment in terms of such parameters as pH and salt concentrations. We hypothesize that enzymes will experience unique chemical environments when incorporated into axonemal protein arrays, and will be most effective in these arrays if the enzyme is selected so that the chemical environment of the flagellum or axoneme is matched to the optimal conditions for catalysis by the enzyme. Microtubules are negatively charged, with a charge density on the microtubule surface of -2.5 e/nm along a protofilament (Minoura et al., 2006; Janmey et al., 2014). FAP20 is located within 10 nm of this charged microtubule surface (Yanagisawa et al., 2014), well within the Debye length for sodium ions at physiological salt (70 nm; Lin et al., 2009) and will, therefore, be within an appropriate distance to feel the altered ionic concentration. We therefore hypothesize that an enzyme docked on the axoneme using FAP20, will be surrounded by a different ionic environment than that of the bulk solution, even in isolated axonemes without a membrane. For enzymes docked using the RSP3 as a scaffold, the effect will be even larger since the radial spoke is buried within the interior of the axoneme. It is interesting to consider whether the distinct charged environments with an axoneme could be exploited.

### Towards solid-phase synthetic biology

In addition to the potential advantages of protein arrays for enzyme applications discussed above, the ability to dock proteins of interest into spatially patterned arrays may open up a new strategy for implementing logic and computational processes using biological molecules. Much work in synthetic biology focuses on solution phase biochemistry. Examples include re-engineering gene regulatory networks in which soluble transcription factors bind and unbind from promoters or re-wiring kinase/phosphatase signaling networks mediated by reversible protein-protein interactions. From a circuit design perspective, the fact that a given protein is free to diffuse throughout the cell means that one protein can only carry one signal. If we want to add a second signal into the circuit, we need to add a second protein. This creates a huge challenge for engineering any but the most trivial of circuits. In electronics, complex circuitry was made possible by the fact that each component is irreversible locked into position, and the interactions between components are rigidly defined by the wires to which they are attached. By locking proteins into position on the axoneme or other insoluble cellular structures, we can limit their potential interactions to their nearest neighbors. How to exploit such spatially restricted interactions is still an open question, but we note that there are already paradigms in computer science and engineering in which an array of identical devices, that can only interact with their immediate neighbors, can carry out complex computational tasks. Specific examples include the large class of models known as “cellular automata” (Toffoli, 1977) as well as fixed networks of locally coupled oscillators (Csaba and Porod, 2020; Mallick et al., 2020)

### Concluding Remarks

We have shown that the flagellar axoneme provides a platform for bio-assembly of linear protein arrays. We find that different axonemal proteins can be used as scaffolds to attach other protein moieties as fusion constructs. Enzymes linked to the axoneme in this way retain activity, but can be liberated from the axoneme using site specific proteases. The axoneme system is highly scalable at low cost given the biological self-assembly of immobilized arrays, the fact that algal cells can be grown without providing a carbon source in the media, and the ability to isolate the arrays using a one-step purification that retains biomass of the producing cell. The uniform, high packing density of axonemal proteins with precise relative protein orientations creates new potential opportunities for substrate channeling, biosensing, and protein-based computation.

## Methods

### Chlamydomonas culture and imaging

*Chlamydomonas* cultures were grown in Tris-acetate-phosphate (TAP) media at 25°C, with constant aeration and continuous light. For imaging fixed cells, *Chlamydomonas* cells were attached to polylysine-coated coverslips and then fixed with 4% paraformaldehyde in PBS for 5 min. Cells were then mounted in Vectashield (Vector Laboratories, Burlingame, CA) on a slide and imaged using a spinning disk confocal microscope (Eclipse Ti; Nikon, Tokyo, Japan) with a 100x oil objective (Apo TIRF, NA 1.49; Nikon), confocal system (CSU-22; Yokogawa Electric Corporation, Japan), and an EMCCD camera (Evolve Delta; Photometrics, Tucson, AZ). 3D images were taken with a z-axis spacing of 0.2 μm.

Flagellar length and fluorescent intensity were measured using custom-written routines in MATLAB (MathWorks, Natick, MA). Z-stack images were convolved with a three-dimensional Gaussian filter to detect the cell body position by finding the center of mass. The cell body was subtracted from the original z-stack images, such that the remaining intensity was restricted to the flagella, after which the position of flagellar tips was detected. Flagellar position was then identified by moving backwards from the flagellar tip step by step on the basis of the local intensity.

### Preparation and storage of flagella and axonemes

Isolation of flagella were performed as described (Richey and Qin, 2013). Flagella were released from the cell body using the pH shock method. Isolated flagella were demembranated by treated with 1% NP-40. Isolated flagella and axonemes were flash frozen in liquid nitrogen and subsequently stored at -80°C.

### TEV protease cleavage

Isolated axonemes were treated with TEV protease (GenScript, Piscataway, NJ) overnight at 4°C, then centrifuged at 20,000 xg for 20 min to pellet the axonemes. The protease-treated axonemes were then attached to poly-L-lysine-coated coverslips, fixed with ice-cold methanol, blocked with 5% BSA, 1% fish gelatin, and 10% normal goat serum in PBS, and then incubated with mouse anti-GFP monoclonal antibody(1:1000; Roche, Indianapolis, IN) as well as rabbit anti-alpha-tubulin polyclonal antibody (1:1000; Abcam, Cambridge, United Kingdom). Following primary staining, axonemes were washed with PBS and incubated with mouse-Alexa488 and rabbit-Alexa546 antibodies (1:200; Invitrogen, Waltham, MA). Samples were washed with PBS and mounted with Vectashield, then imaged using a DeltaVision microscope (GE Healthcare, Chicago, IL) equipped with a 100x objective (Olympus, Tokyo, Japan). Z-stacks were collected at an interval of 0.2 μm, then deconvolved and projected with DeltaVision software (GE Healthcare).

### Fusion constructs

The rescue constructs were modified by substituting the *GFP* gene with the target protein gene in both pBS-FAP20-GFP and pKF-RSP3-GFP. First, the original *Eco*RV linker and *GFP* gene were deleted. In the pKF-RSP3-GFP construct, the C-terminal 140 amino acids of RSP3 were also deleted to generate more space for the target protein because the last 140 amino acids of RSP3 do not affect either recruitment of radial spoke proteins assembled on the axoneme or the motility of flagella (Diener et al., 1993). Furthermore, the target protein genes following the TEV linker were inserted behind the adaptor protein gene.

### Rescue of im5 and pf14

The plasmid FAP20-TEV-βLac was used to rescue the *im5* strain. The plasmid RSP3-TEV-βLac was used to rescue the *pf14* strain. Both plasmids were linearized with *Ssp*I enzyme, linearized FAP20-TEV-βLac are transformed together with the paromomycin-resistant pSI103 plasmid for selection. Plasmid RSP3-TEV-βLac has paromomycin-resistant gene itself. The electroporation technique was applied to transform *Chlamydomonas* cells. Briefly, *im5* or *pf14* cells were cultured in 100 ml TAP medium to dark green, then transferred to 1 L TAP medium with constant aeration. The cells are ready to be transformed when the concentration reaches 106 cells/ml about 20 hours later. Totally, 108 cells were collected and resuspended in 1 ml TAP liquid media containing 60 mM sorbitol. 300 µl cells’ suspension was transferred to a 4-mm electroporation cuvette. For the *im5* cells, 600 ng linearized FAP20-TEV-βLac together with 300 ng pSI103 plasmids were added and mixed evenly. For the *pf14* cells, 600 ng linearized RSP3-TEV-βLac plasmids were used instead. The mixture of plasmids and cells in the cuvettes were chilled on ice for 5 min and then electroporated with an ECM630 electroporator (BTX, Holliston, MA) with the setup parameters of capacitance 50 μF, resistance 650 Ω, and the voltage 825 V. The cells were recovered on the ice for 15 min after electroporation and then transferred to 10 ml TAP medium with 60 mM sorbitol and culture 24 hours in a low light environment for furthermore recovery. The next day, cells were collected and plated on TAP plates containing 10 ng/μL paromomycin to grow 4-5 days to obtain transformants.

### Prescreening rescue colonies by observing cells’ motility and flagellar assembly

After 4- or 5-days’ culturing, colonies resistant to paromomycin antibiotic should be visible on the plate. Transfer all colonies on the TAP plate to 96-well plates with liquid TAP medium. Observe cells’ motility under an inverted microscope (CKX53, Olympus) after 24 hours of culture. In order to further ensure the cells’ motility and flagellar assembly, motile cells from the last step were cultured in 3 ml TAP medium in a test tube for two days, then observe the cells’ motility with an Axioplan phase contrast microscope (Carl Zeiss, Oberkochen, Germany) and 4x objective (Olympus).

### Flagella regeneration

The protocol of flagella regeneration is modified slightly upon the flagella isolation method. Instead of underlay with 25% sucrose immediately after pH shock, the flagella and cell bodies were separated by centrifuging at 2,000 rpm, 5 min, 18°C. The supernatant is containing the flagella and subjected to the 25% sucrose underlay to collect flagella, while the cell bodies were resuspended in 500 ml HEPES to culture 1-2 hours to regenerate flagella. Repeat this repeat if needed to regenerate flagella for more than one time.

### Flagella enzymatic activity assay under different treatments

Soluble βLac enzyme provided in the kit, isolated FAP20-βLac, or RSP3-βLac flagella were freeze-thawed 4 times between -20 and 20°C, treated 10 min at 42°C, or kept at room temperature for 48 hours (overnight). 2.5 μg of commercial soluble βLac enzyme, and 8 μg flagellar samples were subjected to beta-lactamase activity assay. The untreated fresh flagella or soluble commercial protein were used as controls.βLac activity was expressed as nmoles/min hydrolyzed nitrocefin generated per μg of protein (mU/μg).

The protein amount of each sample per reaction was calculated as bellow:

- Commercial βLac: Recombinant protein equals 0.5 μg/μl, added 5 μl per reaction, total protein amount equals 0.5 μg/μl x 5 μl = 2.5 μg.
- WT flagella: 1.6 μg/μl x 5 μl = 8 μg.
- FAP20-βLac flagella: First, ratio of FAP20 and α-tubulin numbers were calculated in the 96 nm repetitive axoneme structure. FAP20 repeat every 8 nm and distributed along 9 doublet microtubules, therefore number of FAP20 equals (96/8) x 9 = 108. The number of βLac equals FAP20, which is 108. α, β-tubulin dimers are 8 nm long and compose 9 doublet microtubules and a center pair of singlet microtubules. The number of α, β-tubulin dimer equals (96/8) x [(13+9) x 9 + 13×2] = 2688. Second, because the molecular weight of FAP20-βLac is 40 kDa and molecular weight of a dimer is 100 kDa, the mass ratio of βLac and α, β-tubulin dimer equals (2688/108) x (100/40) = 62. Finally, the mass of βLac in the FAP20-βLac flagella sample equals (1.6 μg/μl x 5 μl)/62 = 0.13 μg.
- RSP3-βLac flagella: First, the ratio of RSP3 and α-tubulin numbers were calculated in the 96 nm repetitive axoneme structure. Radial spoke has an average number of 2.5 every 96 nm and forms a dimer, therefore the number of RSP3 per flagellum is 2.5 × 2 × 9 = 45. The number of βLac equals RSP3, which is 45. The number of α, β-tubulin dimer is 2688. Second, because the molecular weight of RSP3-βLac is around 100 kDa and molecular weight of α, β-tubulin dimer is around 100 kDa, the mass ratio of βLac and α-tubulin equals (2688/45) x (100/100) = 60. Finally, the mass of βLac in the RSP3-βLac flagella sample equals (1.6 μg/μl x 5 μl)/60 = 0.13 μg.

Once the commercial βLac enzyme or flagella samples mixed with nitrocefin to start the reactions, the absorbance of 490 nm for each reactant was measured every minute until the absorbance reached plateau. The concentration of the products was calculated based on the standard formula y = 0.0185x + 0.0318 (**Supplemental Figure S2**). The enzymatic activity was calculated at 1 min, 2 min, and 3 min. The enzymatic activity was calculated by averaging three measurements with standard deviation.

## Acknowledgments

We thank members of the Marshall and Qin laboratories for helpful discussion. This work was supported by NIH grant R35 GM130327(W.F. M.) and by the Center for Cellular Construction, an NSF-funded Science and Technology Center supported by NSF grant DBI1548297 (W.F.M.)

